# Sex differences in post-stroke aphasia rates are caused by age. A meta-analysis and database query

**DOI:** 10.1101/407296

**Authors:** Mikkel Wallentin

## Abstract

**Background:** Studies have suggested that aphasia rates are different in men and women following stroke. One hypothesis says that men have more lateralized language function than women. Given unilateral stroke, this would lead to a prediction of men having higher aphasia rates than women. Another line of observations suggest that women are more severely affected by stroke, which could lead to a higher aphasia rate among women. An additional potential confounding variable could be age, given that women are typically older at the time of stroke.

**Methods & Procedures:** This study consists of two parts. First, a meta-analysis of the available reports of aphasia rates in the two sexes was conducted. A comprehensive literature search yielded 25 studies with sufficient information about both aphasia and gender. These studies included a total of 48,362 stroke patients for which aphasia rates were calculated. Second, data were extracted from an American health database (with 1,967,038 stroke patients), in order to include age and stroke severity into a regression analysis of sex differences in aphasia rates.

**Outcomes & Results:** Both analyses revealed significantly larger aphasia rates in women than in men (1.1-1.14 ratio). This speaks against the idea that men should be more lateralized in their language function. When age and stroke severity were included as covariates, sex failed to explain any aphasia rate sex difference above and beyond that which is explained by age differences at time of stroke.

## Introduction

A stroke is a medical condition in which blood flow to the brain is restricted, due to occlusion (ischemic stroke) or hemorrhage (hemorrhagic stroke), resulting in cell death (WHO). In the US alone, approximately 800,000 people experience a stroke every year, according to the American Heart Association [1]. Stroke is the leading cause of motor and cognitive disability in western countries and aphasia, the inability to comprehend and formulate language because of brain damage, is one of the most common deficits after stroke. A large variability in the reported frequency of aphasia can be found in the literature [e.g. 2, 3–5], ranging from 15 % to 68 % of acute patients. A recent meta-analysis, however, concluded that aphasia is present in approximately 30 % of acute patients and 34 % in rehabilitation settings [6]. Variability of measured rate of aphasia has many causes. The method for aphasia identification differs between hospitals and countries and some sub-scores for aphasia in stroke scales have been found to be limited in their accuracy and reliability [7, 8]. Another potential source of variance may be sex. The above-mentioned meta-analysis did not take potential sex differences into account.

Stroke has been noted to affect the sexes differently. Stroke has been reported to be more common among men [9]. The symptoms of stroke have also been found to differ somewhat between men and women. Women are often more severely affected overall, more often experience paralysis, impaired consciousness and altered mental status together with a generalized weakness, while men more often experience dysarthria, diplopia, sensory loss, ataxia and balancing problems [10]. An association between pre-stroke dementia, which is more prevalent in women, and stroke severity has also been noted [11]. Lastly, aphasia following stroke has been reported to affect women to a larger degree than men [see 10 for a review], although evidence has been conflicting [e.g. 12, 13, 14].

Potential sex differences in aphasia may shed light on overall sex differences in language and cognition. Sex differences in certain linguistic domains are known to exist within the normal population, with differences in first language acquisition speed [15] and reading and writing abilities [16] being the most consistent, favoring girls/women over boys/men. Differences in word use have also been documented [17]. The underlying causes for these differences are probably complex and research trying to tie them to brain structure and function has yielded inconsistent results [18]. Some studies, however, have argued for the hypothesis that language is more bilaterally organized in the brains of women compared to men [e.g. 19, 20–22], although this is highly controversial [18, 23, 24]. A sex difference in language lateralization would ultimately lead to a sex difference in aphasia following unilateral stroke. If men’s language is more lateralized in the brain than the language of women, we would expect them to be more prone to aphasia following unilateral stroke and vice versa, if women have greater language lateralization than men, we would expect women’s language function to be more vulnerable to stroke.

In this paper I conduct a meta-analysis on aphasia rate given stroke across published peer-reviewed papers and test if the frequency for the two sexes differ. I then compare the results of the meta-analysis to data from a large American patient database (Healthcare Cost and Utilization Project (HCUP) under Agency for Healthcare Research and Quality, U.S. Department of Health & Human Services: https://hcupnet.ahrq.gov). Given that both analyses are based on fully anonymized and publicly available data, the study poses no ethical concerns.

## Meta-analysis methods

This report was prepared according to the Preferred Reporting Items for Systematic Reviews and Meta-Analyses (PRISMA, http://www.prisma-statement.org). PRISMA is an evidence-based minimum set of items for reporting in systematic reviews and meta-analyses [25]. The supporting PRISMA checklist for this meta-analysis is available as supporting information; see S1 PRISMA Checklist.

The main outcome measures for the analysis were aphasia rate (percentage of stroke patients with aphasia diagnosis) and the sex ratio of aphasia rates. A pub-med search including the terms “stroke” AND “aphasia” AND “gender” (which automatically includes the term “sex”) generated 211 citations up until July 1st 2018 (see Fig 1 for a flow-chart of the data sampling procedure). No time or language constraints were put on the sampled reports. References in review articles on stroke and aphasia were also investigated. A total of 419 titles were considered. 91 papers were selected for further inspection on the basis of their title and abstract. The analysis set out to study post-stroke aphasia in the broadest sense. Speech pathology tests still lack standardization and diagnostic data for identifying aphasia in stroke populations [26]. Different diagnostic approaches to aphasia were therefore not distinguished. All studies including some overall aphasia diagnosis were deemed relevant. Inclusion criteria were the following: 1) Primary peer reviewed studies reporting aphasia frequency among stroke patients; 2) Studies reporting overall number of aphasia patients and stroke patients from a unselected stroke cohort (thus limiting bias), i.e. studies dealing only with sub-types of aphasia were excluded and studies of group comparisons between aphasia patients and matched control groups were also not considered; 3) Studies reporting aphasia counts for both male and female aphasia patients as well as for stroke patients from both sexes or reports where these numbers could be extracted from reported percentages; 4) First/Primary report of data: The same data could only be included once. If more than one paper investigated the same patient group, the earliest/most comprehensive publication was chosen.

**Fig 1.**
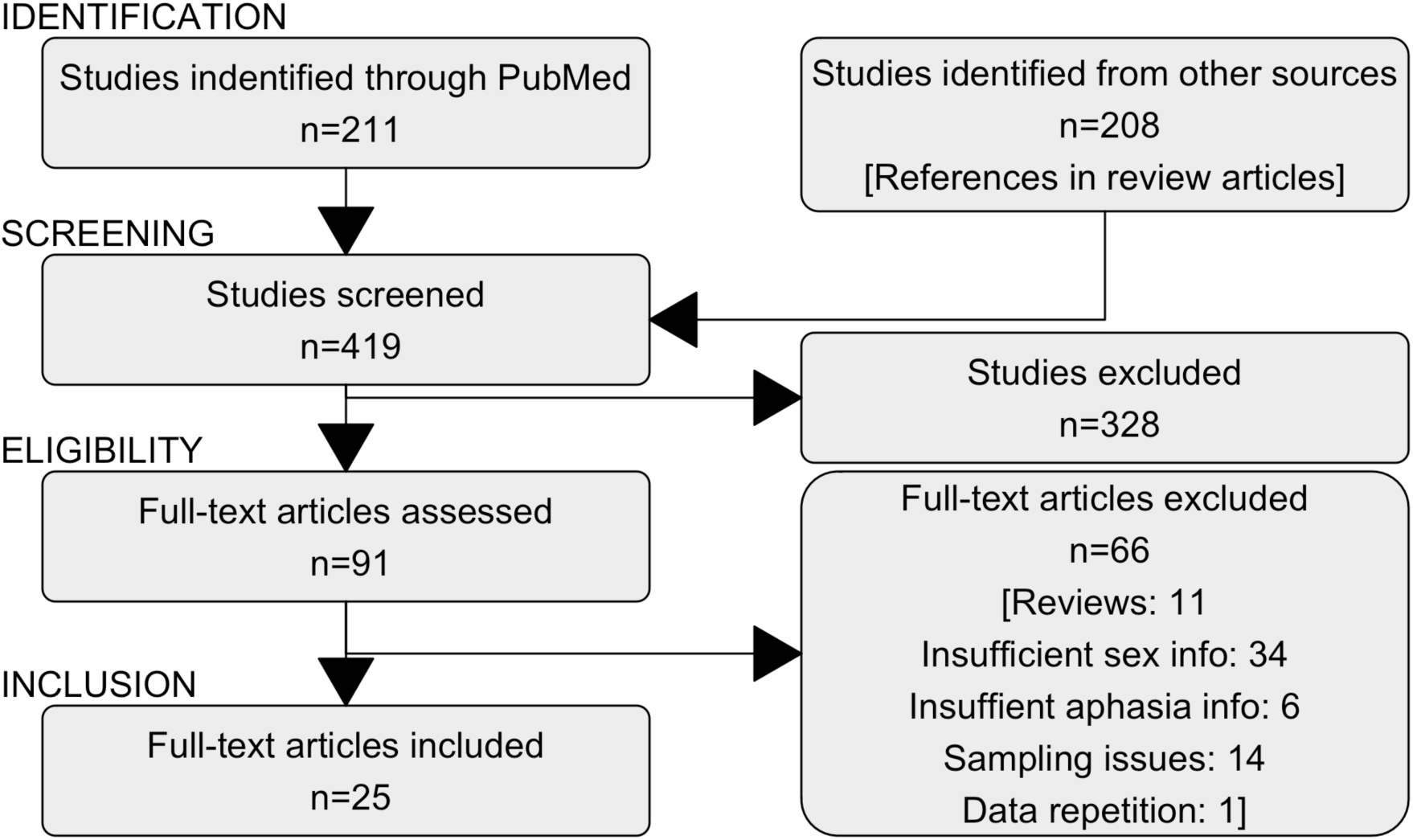
Data gathering flow-chart. Flow chart depicting the different phases of data gathering.

Twenty-five studies were included in the final dataset [4, 5, 11–14, 27–45]. One study [12] contained information about aphasia frequency for different age bands. These groups were included separately (see Table 1 and Fig 2). I excluded 11 reviews [6, 9, 10, 46–53], 34 studies with insufficient information about the sexes [2, 54–86], 6 studies with insufficient aphasia information [87–92], 14 studies with insufficiently specified sampling procedure for aphasia patients from stroke cohorts [93–106], and one study with repetition of data use [38] (see Fig 1).

**Table 1.** Studies included in the meta-analysis by publication date.

| Study | Place | Note | N (stroke) | Aphasia Rate<br>% Female | Aphasia Rate<br>% Male | Ratio | 95% CI<br>lower | 95% CI<br>upper |
| --- | --- | --- | --- | --- | --- | --- | --- | --- |
| Brust et al. 1976 [32] | New York, USA | acute stroke | 850 | 18.4 | 23.8 | 0.774 | 0.580 | 1.040 |
| Siirtola et al. 1977 [45] | Turku, Finland | acute stroke | 338 | 28.7 | 28.2 | 1.016 | 0.670 | 1.540 |
| Miceli et al. 1981 [14] | Rome, Italy | 223 CVA, 128 tumors, 29 traumas etc. | 390 | 60.8 | 62.3 | 0.975 | 0.750 | 1.280 |
| Scarpa et al. 1987 [44] | Modena, Italy | left hemisphere stroke, post 14 days | 196 | 62.5 | 50.0 | 1.250 | 0.860 | 1.820 |
| Hier et al. 1994 [33] | four sites, USA | acute stroke | 1805 | 22.6 | 19.4 | 1.103 | 0.920 | 1.330 |
| Pedersen et al. 1995 [13] | Copenhagen, Denmark | acute stroke | 881 | 39.7 | 34.8 | 1.140 | 0.920 | 1.420 |
| Laska et al. 2001 [29] | Danderyd, Sweden | acute stroke | 106 | 33.3 | 34.7 | 0.961 | 0.500 | 1.850 |
| Godefroy et al. 2002 [5] | Lille, France | Acute stroke, Aphasia | 308 | 68.5 | 66.1 | 1.037 | 0.790 | 1.360 |
| Di Carlo et al. 2003 [39] | 7 European countries | acute stroke | 4499 | 34.8 | 30.3 | 1.337 | 1.060 | 1.690 |
| Kelly-Hayes et al. 2003 [87] | Framingham MA, USA | at 6 months post stroke | 108 | 23.8 | 11.6 | 2.143 | 0.780 | 5.900 |
| Roquer et al. 2003 [30] | Barcelona, Spain | acute stroke | 1581 | 28.9 | 21.6 | 1.335 | 1.100 | 1.630 |
| Engelter et al. 2006 [34] | Basel, Switzerland | acute ischemic stroke | 269 | 34.0 | 23.9 | 1.422 | 0.890 | 2.260 |
| Kyrozis et al. 2009 [28] | Acadia, Greece | 28 days post-stroke | 555 | 27.6 | 18.8 | 1.473 | 1.040 | 2.090 |
| Tsouli et al. 2009 [4] | Athens, Greece | acute stroke | 2297 | 41.3 | 31.5 | 1.313 | 1.140 | 1.510 |
| Brkic et al. 2009 [31] | Tuzla, Bosnia and Herzegovina | acute stroke | 993 | 23.0 | 17.6 | 1.309 | 0.990 | 1.730 |
| Bersano et al. 2009a [12], <64 years | seven regions, Italy | acute stroke, <64 years | 1751 | 21.0 | 20.0 | 1.044 | 0.840 | 1.300 |
| Bersano et al. 2009b [12], 64-74 years | seven regions, Italy | acute stroke, 64-74 years | 2663 | 26.0 | 24.0 | 1.081 | 0.930 | 1.260 |
| Bersano et al. 2009c [12], 75-84 years | seven regions, Italy | acute stroke, 75-84 years | 2853 | 31.0 | 27.0 | 1.160 | 1.010 | 1.330 |
| Bersano et al. 2009d [12], >84 years | seven regions, Italy | acute stroke, 84< years | 1581 | 43.0 | 35.0 | 1.223 | 1.030 | 1.450 |
| Dickey et al. 2010 [36] | Ontario, Canada | at discharge | 15327 | 33.4 | 31.0 | 1.078 | 1.020 | 1.140 |
| Gall et al. 2010 [11] | Melbourne, Australia | acute stroke, Dysphasia | 843 | 46.1 | 35.6 | 1.294 | 1.040 | 1.600 |
| Gialanella et al. 2011 [38] | Lumezzane, Italy | acute stroke | 262 | 55.9 | 44.4 | 1.258 | 0.890 | 1.770 |
| Croquelois & Bogousslavsky 2011 [27] | Lausanne, Switzerland | acute stroke | 5880 | 28.1 | 24.9 | 1.128 | 1.020 | 1.250 |
| Jerath et al. 2011 [41] | Rochester, USA | acute stroke | 449 | 45.8 | 37.4 | 1.222 | 0.910 | 1.640 |
| Kadojic et al. 2012 [37] | Osijek, Croatia | acute ischemic stroke | 177 | 48.2 | 37.2 | 1.294 | 0.820 | 2.040 |
| Flowers et al. 2013 [35] | Toronto, Canada | acute ischemic stroke | 221 | 35.7 | 26.0 | 1.373 | 0.850 | 2.220 |
| Wasserman et al. 2015 [42] | Ottawa, Canada | isolated aphasia as only deficit of stroke | 1155 | 4.1 | 3.0 | 1.347 | 0.720 | 2.510 |
| Chang et al. 2015 [43] | Colombo, Sri Lanka | data from questionnaire | 24 | 62.5 | 56.2 | 1.111 | 0.370 | 3.320 |
| **ALL** |  |  | **48362** | **29.6** | **26.0** | **1.139** | **1.100** | **1.180** |

Overview of the studies included in the meta-analysis on sex differences in post-stroke
aphasia rate.

**Fig 2.**
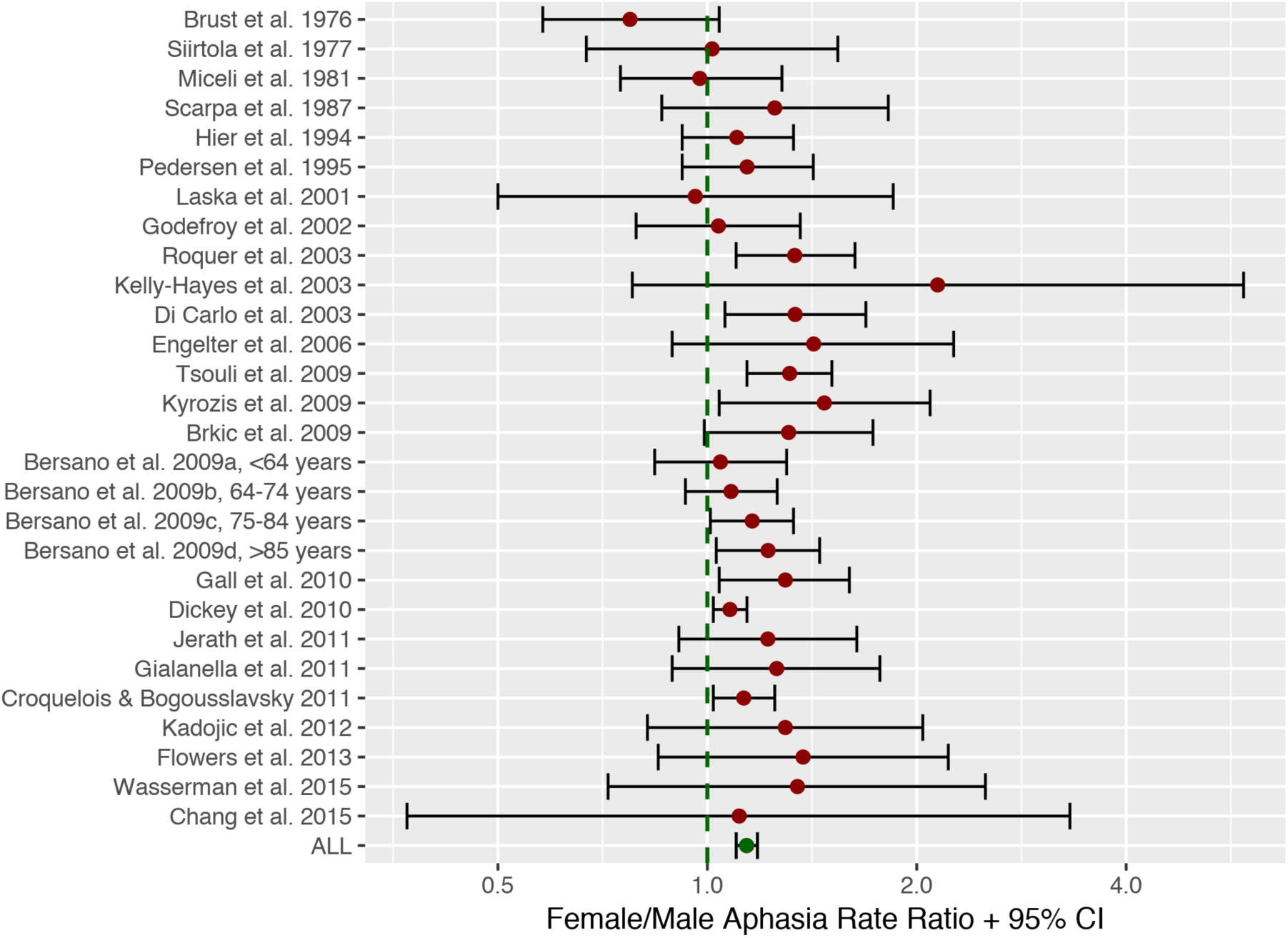
Meta-analysis forest plot. Forest plot of aphasia rate ratios between males and females for the 25 studies included in the meta-analysis (total n=48,362), showing that across studies a small but significant effect of sex exists, indicating that women are more likely to get aphasia from stroke. This effect, however, does not take age or stroke severity into account.

For every study, the number of stroke patients for each sex was extracted together with the number of aphasia patients for each sex and the percentage of aphasia patients. Studies with more patients were considered less at risk of within-study bias and analyses therefore incorporated study size weights. Study year, and place were also noted, as well as potential additional information about the patients’ age and time/type of study relative to disease onset. All data were analyzed using R. Heterogeneity between studies was assessed using the metaphor package [107] and a funnel plot was used to inspect potential publication/selection bias. Year of study was added as a covariate to a subsequent analysis in order to investigate if sex differences in aphasia diagnoses have been changing over time.

## Results of meta-analysis

The 25 studies included a total of 48,362 stroke patients (23,085 women, 25,297 men). Of these 13,398 (6,828 women, 6,570 men) were diagnosed with aphasia (27.7%). 29.6 % of female stroke patients were diagnosed, while 26 % of males were diagnosed with aphasia (see Table 1 and Fig 2). This difference was found to be statistically significant using a paired and weighted t-test on the aphasia rates across studies, weighted to add emphasis on studies with larger patient samples, *t*(27)=6.76, *p*<0.001, forcing a rejection of the null-hypothesis that there is no difference in aphasia rate between women and men. The overall sex aphasia rate ratio was found to be 1.14 (1.10-1.18 95% CI) with a Cohen’s *d* of 0.37 which is usually considered a small effect [108]. Low to medium heterogeneity between studies was observed: *I*^2^= 41 % (CI: 4%-77%) (see Fig 1 for a forest plot) and a funnel plot did not suggest any outspoken bias in the reports (see Fig 3). Including publication year as a covariate in a regression analysis did not alter the result and was not in itself a significant predictor of sex differences (p>0.05).

**Fig 3.**
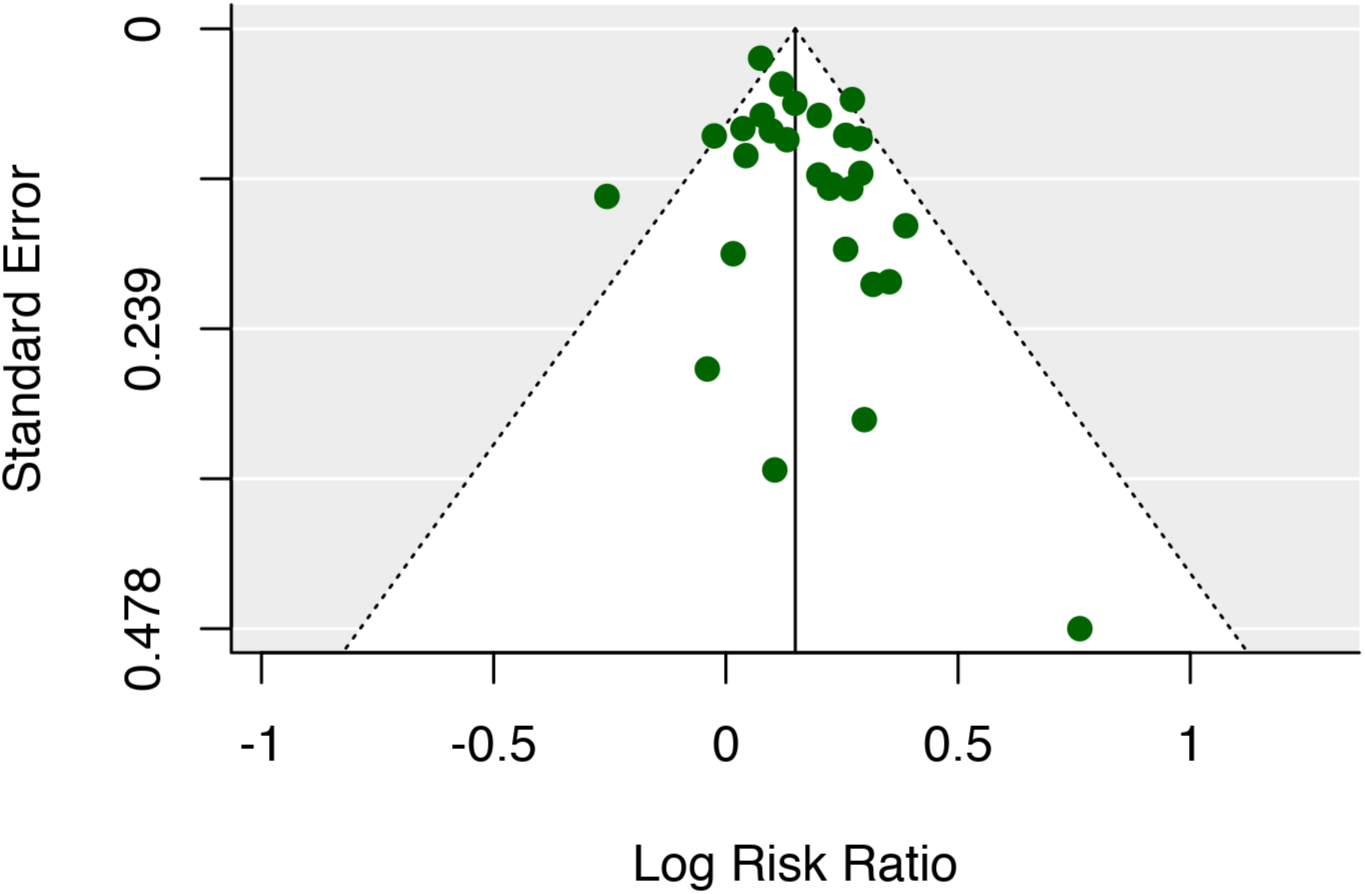
Funnel plot. A Funnel plot did not indicate any outspoken bias in the meta-analysis data.

## Interim discussion

A higher aphasia rate after stroke for women than for men was found across studies in the meta-analysis. The average aphasia rate for women was 29.6% and for men 26% (see Table 1). The effect-size was in the small range.

The aphasia rate across studies and sexes (27.7%) was comparable to that reported in a recent meta-analysis (30%) [6]. The slightly lower estimate may in part be related to the inclusion of a study of cases with isolated aphasia [42] which had a much smaller aphasia rate than studies with a regular aphasia diagnosis (see Table 1). The aphasia rate ratio for the sexes reported on isolated aphasia by [42], however, was comparable to that of most other studies in the sample. Study year did not explain any variance in sex differences, suggesting that e.g. evolving diagnostic procedures for aphasia [26] have not had an observable linear impact on sex differences in aphasia rates. The forest plot in Fig 2 is ordered according to publication year on the y-axis, and given that there had been changes as a function of time, this should be visible as either a leftward or rightward trend in the data points. This seems not to be the case.

The fact that a higher aphasia rate after stroke for women than for men was found across studies contradicts the notion that language in men is more lateralized than in women (see introduction). If men have more lateralized language, one would expect their language to be more vulnerable to unilateral stroke than women’s language, which, as shown, is not the case. But the findings are also at odds with previous critical suggestions that there are no sex differences in language lateralization between women and men [e.g. see 18, 109]. At face value, the findings would suggest that women in fact have more lateralized language than men. There are, however, reasons to hesitate before arriving at such a conclusion, based on the present analysis. As mentioned in the introduction, stroke is known to affect men and women differently on a number of accounts, including general severity. The sexes also differ on general health levels, resulting in women being older, on average, when they get a stroke [9]. Age has previously been found to be a predictor of acquiring aphasia [52]. In order to investigate if the effects of sex found in the meta-analysis are specific to language or may relate to more general differences that are unlikely to be caused by a sex difference in core language function, an investigation of aphasia rates that include additional explanatory variables is needed. Unfortunately, very few studies in the current cohort make detailed reports of age effects on aphasia stratified for sex. One exception is Bersano et al. [12] who report aphasia rates for 4 different age groups. Here an interaction between age and sex differences seemingly can be observed. The sex difference is almost non-existing in the youngest age group (under 64), but gradually grows larger and larger in older age groups (see Table 1). It thus seems that taking age into account is important when trying to understand the sex difference in aphasia rates.

Only six studies provided precise gender-stratified age information for their samples (see Table 2). The female stroke groups are older than the male groups in five of these six studies. A post-hoc weighted t-test again found that females were more likely to suffer from aphasia, *t*(5)=5.1, *p*<0.005, but when adding age difference as a covariate, this effect was no longer significant, *t*(4)=0.8, *p*>0.05. It could thus seem that age explains some of the difference found between the sexes. The low number of degrees of freedom and the diverse study characteristics, however, make it very difficult to conclude anything definite on the basis of this analysis.

**Table 2.** Studies in the meta-analysis with age information.

| Study | N (stroke) | Aphasia Rate<br>% Female | Aphasia Rate<br>% Male | Ratio | Age<br>Female | Age<br>Male | Age<br>difference |
| --- | --- | --- | --- | --- | --- | --- | --- |
| Hier et al. 1994 [33] | 1805 | 22.6 | 19.4 | 1.165 | 69.2 | 65.3 | 3.9 |
| Kelly-Hayes et al. 2003 [87] | 108 | 23.8 | 11.6 | 2.052 | 80.3 | 75.8 | 4.5 |
| Roquer et al. 2003 [30] | 1581 | 28.9 | 21.6 | 1.335 | 74.6 | 68.8 | 5.8 |
| Gall et al. 2010 | 843 | 46.1 | 35.6 | 1.294 | 76.0 | 72.0 | 4.0 |
| Jerath et al. 2011 [41] | 449 | 45.8 | 37.4 | 1.225 | 79.0 | 70.0 | 9.0 |
| Chang et al. 2015 [43] | 24 | 62.5 | 56.2 | 1.111 | 61.6 | 64.7 | -3.1 |

Another possible explanation for the higher aphasia rate in women is that they may be affected more severely by stroke in a non-discriminant manner [10]. If aphasia rates can be explained by severity alone, it would again suggest that the sex difference is not restricted or related to language in any meaningful way. But again, this type of information is not reported in the papers covered by the present meta-analysis.

An additional limitation in the present meta-analysis is that the literature has been surveyed and studies have been selected by a single individual, the author. This carries the potential risk of conscious or nonconscious selection bias. Speaking against this is the fact that the observed sex difference in aphasia goes directly against the conclusions of the author’s prior work. I have argued against the existence of large sex differences in the brain architecture supporting language, both in review articles [18, 110] and based on neuroimaging studies [109, 111]. Before starting the data collection, I assumed that there would be no differences in aphasia related to sex. If anything, a potential selection bias based on the author’s bias, would most likely mean that the observed sex difference is underestimating the real difference. Inspection of the funnel-plot did not reveal bias in the selected studies, and I consider this risk rather small, but it is a possibility.

One way to confront this problem and counter the potential age and severity confounds at the same time is to get access to more detailed aphasia data with additional age and severity information. One suggestion could be to contact the authors of the studies included in the meta-analysis and try to obtain the relevant information. However, given the large time span covered by the included publications, this would be unfeasible. I have therefore instead added a 2^nd^ dataset from an American healthcare database (see below) that will allow me to investigate aphasia rates while taking age and stroke severity into account.

## Methods for database analysis

Data from the Healthcare Cost and Utilization Project (HCUP) from community hospitals in the United States were used for the analysis. The database (https://hcupnet.ahrq.gov/) contains data from the National Inpatient Sample (NIS) using the International Classification of Diseases and Health Related Problems (ICD-9) codes from 35 US American states from the years 2011-2014. The ICD system is used by US hospitals for reimbursement purposes and subsequent research, e.g. to study patterns and outcome of disease [112]. Starting in data year 2012, the NIS is a sample of discharge records from all HCUP-participating hospitals. For prior years (i.e. including 2011), the NIS was a sample of hospitals from which all discharges were retained. Patient counts for each year from each state, stratified by sex, was used in the analysis. Data from this database has previously been used to study post-stroke aphasia rates [2], but here we add sex, age and severity as explanatory variables and incorporate all available states for all the years in which the ICD-9 diagnoses were used (i.e. 10 times more patients).

To identify number of patients with stroke, the combined number of diagnoses from the database related to stroke was obtained. To emulate the studies in the meta-analysis as much as possible, both hemorrhagic and ischemic stroke cases were included in the analysis. Cases with the following ICD-9 codes were thus selected: “431 Intracerebral Hemorrhage“, “434.00 Crbl Thrmbs Wo Infrct”, “434.01 Crbl Thrmbs W Infrct”, “434.10 Crbl Emblsm Wo Infrct”, “434.11 Crbl Emblsm W Infrct”, “434.90 Crbl Art Oc Nos Wo Infrc”, “434.91 Crbl Art Ocl Nos W Infrc”, “436 Cva”. Cases under the ICD-9 codes “433*” (occlusion and stenosis of precerebral arteries) were not included because they have been found to have poor predictive power for acute stroke [113]. To identify the number of patients with aphasia, the ICD-9 code: “784.3 Aphasia” was used. As a proxy for stroke severity, the number of hemiparesis/hemiplegia diagnoses were included (using the ICD-9 code: “342.90 Unsp Hemiplga Unspf Side”, which is the most commonly used ICD-9 code for hemiplegia/paresis in the database and which does not bias towards the dominant or nondominant side). Hemiparesis, hemiplegia and aphasia are comorbid deficits. Around 90% of aphasia patients have hemiparesis [64], but if a sex difference in number of aphasias is accompanied by a similar sex difference in hemiparesis/hemiplegia diagnoses, then the difference is likely to be explained by stroke severity rather than being a specific language related phenomenon.

The database allows for two different ways to draw data. Either one can draw “Principal” diagnoses or “all-listed” diagnoses. As aphasia is often unlikely to be the principal diagnosis in a hospital visit, “all-listed” diagnoses were used. However, age information is only available with “principal” diagnoses, and age information was therefore drawn from this dataset. The assumption is that age differences in principal diagnosis will be representative for age differences in the “all-listed” diagnoses as well. It is important to note that the database does not offer information about age for any individual patient. These data are all summery data for the participating states for a particular year (2011-2014). Yet, similarly to a meta-analysis of data from different studies, summary data from different states, each representing a large group of patients may nevertheless be representative of generalizable differences between groups (males and females) in the data.

To evaluate statistical significance of the predictors, a linear mixed-effects regression analysis was conducted, fit by REML, using the *lmertest* package in R [114]. P-values were estimated using Satterthwaite's method. The model incorporated aphasia rate as the dependent variable and sex as the main fixed dependent variable. Age and rate of hemiplegia diagnoses (proxy for stroke severity) were z-score scaled and added as additional covariates. The model also included all possible interactions between the three variables. US state and year for each data point were included as random effects. The regression was weighted by number of stroke cases in a particular state/year, to put more weight on data-points from larger states.

## Results of database analysis

A total of 1,967,038 stroke patients were found in the database (1,014,239 women, 952,799 men) in the period from 2011 to 2014. Aphasia was diagnosed in 623,942 cases (336,604 women, 287,338 men) or 31.7%. Using this method, 33.2 % of female stroke patients were diagnosed, while 30.2 % of males were diagnosed with aphasia (see Fig 4).

**Fig 4.**
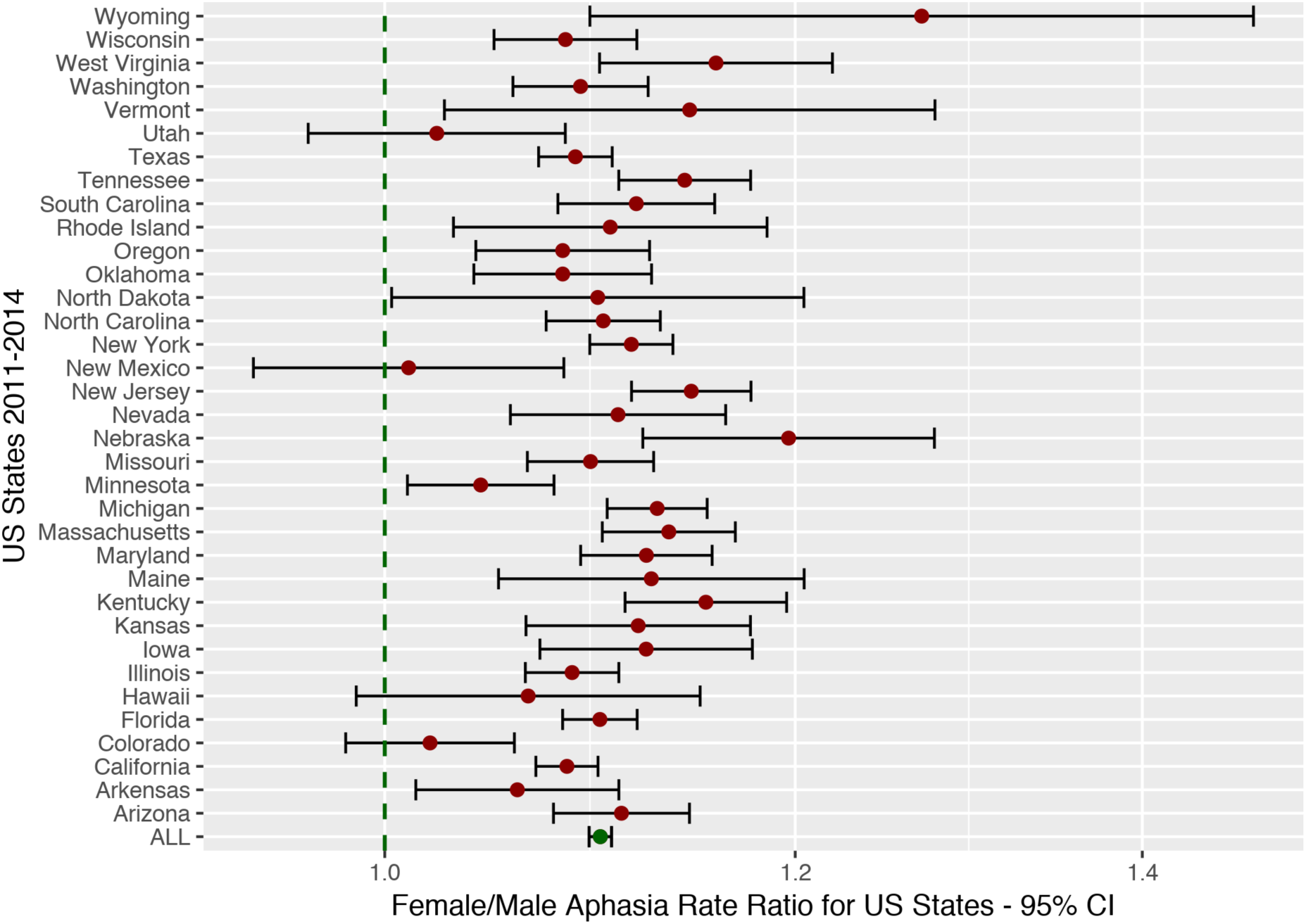
US data forest plot. Aphasia rate ratios (uncorrected for age) for each US state in the HCUP database from 2011-2014. This analysis replicates the findings from the meta-analysis and provides unequivocal evidence for a higher aphasia rate among women compared to men given stroke (see Fig 1, but note the scale difference between plots). However, as Fig 5 shows, this effect can be explained completely by the sex difference in age at stroke.

The overall female/male aphasia rate ratio was found to be 1.100 (1.095-1.106 95% CI) with a Cohen’s *d* effect size across states of 0.63 which is usually considered a medium effect size [108]. A paired t-test again yielded support to the existence of a sex difference, *t*(143)=-18.36, *p*<0.001.

When including age and stroke severity in a regression analysis, however, no significant effect of sex over and above that explained by age and severity could be observed, *t*(273.11)=-1.64, *p*=0.1. A significant effect of age, *t*(274.83)=2.11, *p(uncorrected)*<0.05, and a significant effect of stroke severity, *t*(268.73)=4.77, *p*<0.001 was observed. Fig 5 displays how sex is completely confounded by age of stroke and does not add any explanatory power to the analysis. A significant interaction between age and stroke severity was also observed, *t*(275.31)=-2.02, *p(uncorrected)*<0.05. No other interactions were significant.

**Fig 5.**
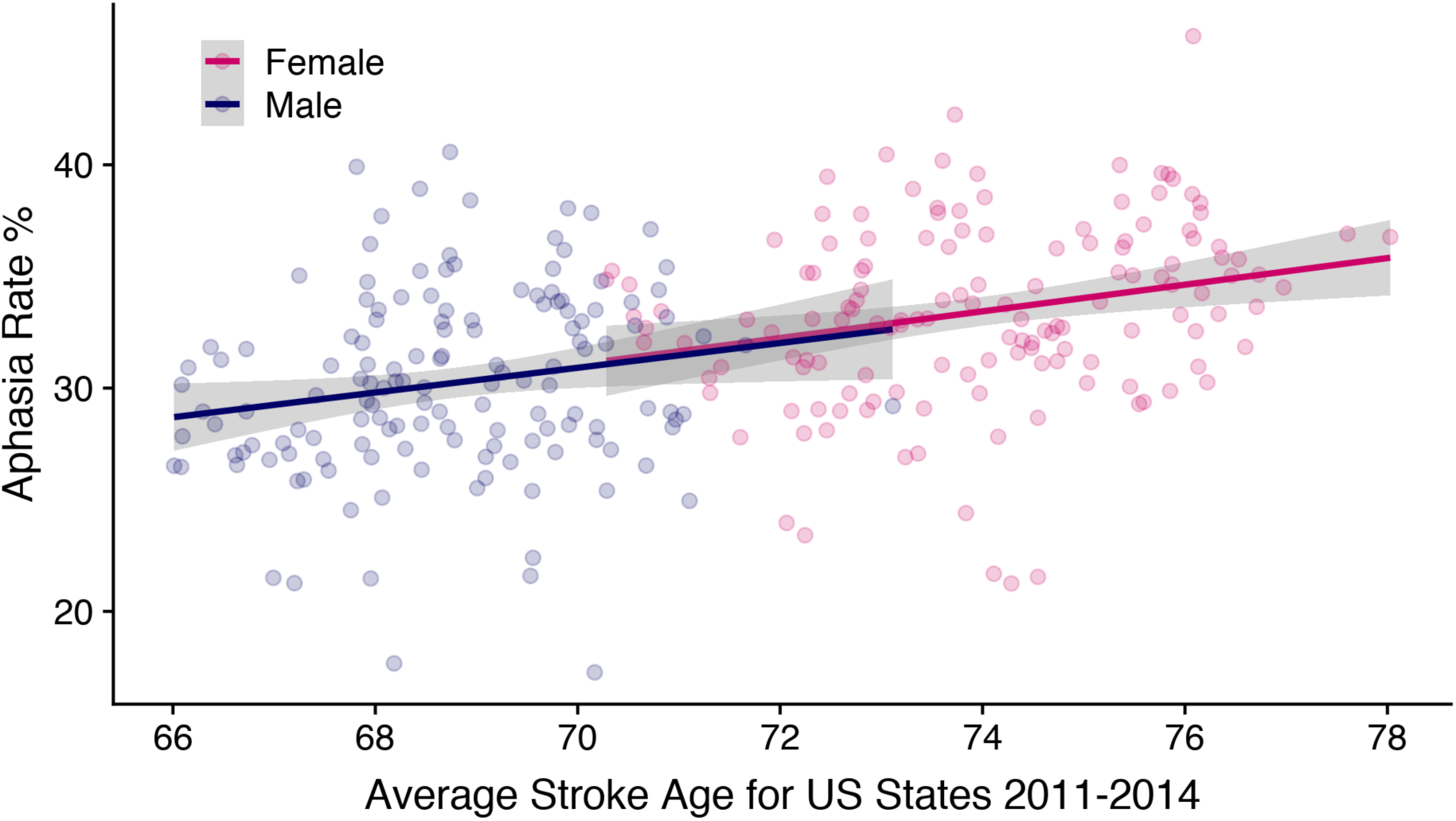
Aphasia rate as a function of age at stroke. A scatterplot of stroke average age against aphasia rate for each US state and year (2011-2014) in the HCUP database. The plot illustrates the large age difference between men and women at time of stroke. It also shows a positive correlation between average age and aphasia rate, suggesting that older stroke patients more often get aphasia. When this relationship is taken into account, sex effects are no longer significant in the aphasia rates.

## Discussion

On this very large cohort of patients, we replicate the findings from the meta-analysis. The overall aphasia-rate for the two studies are compatible. The meta-analysis suggested a weighted average aphasia-rate of 27.7%, whereas the aphasia-rate estimated from the database was 31.7%. Both results are comparable to the estimate reported in another recent meta-analysis (30%) [6], where sex was not considered. Both analyses revealed that, based on raw aphasia rates, women are more likely to get an aphasia diagnosis following stroke than men. The sex aphasia rate ratio was slightly smaller in the database study (1.10) than in the meta-analysis (1.14), but the confidence intervals overlap. This speaks against any large selection bias in the meta-analysis. The effect size, as measured by Cohen’s *d*, was found to be larger in the database-analysis (0.63) than in the meta-analysis (0.37). This is likely due to the fact that the database data is much more homogeneous than the data included in the meta analysis, where the publication time differs substantially (covering the years 1976 to 2015), the time of diagnosis post-stroke differs and the definition of aphasia also differs to some extent (e.g. one study includes dysphasia as well as aphasia and one study only looks at isolated aphasia cases – see Table 1 for details).

At the same time, the 2^nd^ analysis supports the findings from the post-hoc analysis, where age was found to explain away the effect of sex in the subset of papers that had age information. In the database, we find no evidence of any sex difference in aphasia rates over and above that which can be explained by the age differences between females and males at time of stroke. This replicates previous findings that age is a predictor of aphasia following stroke [52], and given that age is a more fundamental causal variable than language (i.e. your language cannot change your age, but the opposite may be true), it is likely that age is the cause of this statistical relationship and also that most, if not all, of the sex differences in aphasia rates are caused by the age difference in stroke between women and men. Figure 5 illustrates this. The regression line for aphasia rate as a function of age for females is an exact continuation of the same regression for males. No offset to imply a main-effect of sex.

I also found an independent effect of stroke severity on aphasia rates as measured by diagnoses of hemiplegia. Aphasia and hemiparesis/hemiplegia are known to be highly comorbid. In this study I found that severity effects on aphasia are independent of the sex effects. The sex differences thus do not seem to be related to stroke severity per se. It has to be said, however, that this analysis used a somewhat crude proxy for stroke severity. Other measures, such as general stroke scale scores [115, 116] might interact more with sex.

Bersano et al. [12] found indications of increasing sex differences in aphasia rates with age. Contrary to this, the database analysis showed no indication of an interaction between sex and age. Bersano and co-workers did not report inferential statistics documenting an actual interaction, but looking at their data, the increasing discrepancy in aphasia between males and females with age is striking. For patients below 64 years the aphasia rate gender ratio is 1.04 and grows to 1.08 in 64 to 74 year-old patients, 1.16 in 75 to 84 and 1.22 in patients above 84 years of age (see Table 1). How does this fit with the current data not showing any interaction between age and sex? One possible explanation is that there is an inherent bias in the way that the Bersano and co-workers’ age data is distributed. When lumping the data into 10-year age bins, one needs to consider that women and men may not be equally distributed within each bin. The data from the Bersano et al. study were collected in 2001 in Italy. If one looks at the gender and age distribution of the Italian population in January 2002 using population statistics (http://demo.istat.it/pop2002/index_e.html), one finds that because women live longer than men, the average age of women within the different age bins from middle age and onwards is higher than that of men and that this difference gets larger for the older groups. For the 54 to 64 year-old Italians, the mean age difference between men and women is 0.07 years, but for the age group above 84 years, it has grown to 0.52 years. There is a very strong linear correlation between the mean age differences in the Italian population in these age bins and the reported differences in aphasia rate (*r*= 0.96, data available from author on request), which suggests that at least some of the interaction between sex and age seen in the Bersano and co-workers’ data is based on unequal sampling of the different ages. This is not to say that there could not be sex and age interactions that were not picked up by the current analysis. The data from the database is distributed on a state by year basis and each data-point for age is the result of averaging across many individual patients. Underneath this gross data reduction may be hidden lots of important variability. Further studies are needed in order to rule out a potential interaction between sex, age and aphasia.

The present analyses are also limited in that they say nothing about the different types of aphasia symptoms that patients may suffer from and the potential interactions that might be found with gender if one looks more carefully at aphasia subtypes.

Taken together, the results are in line with a critical stance towards any large-scale brain base for sex differences in language [18]. This, of course, does not mean that the observed sex differences in language related behavior (see introduction) do not have brain correlates, just that these differences will be dynamic, complex and to a large extent dependent on gender differences in experience and context rather than being tied to genetic sex.

Aphasia is a strong predictor of clinical outcome in stroke [4, 6]. The clinical implications of the present results are that sex can be used as a weak predictor for aphasia in stroke patients in the absence of knowledge about age. This, of course, will only have limited applicability. Apart from this, the findings are in line with results from stroke treatment studies that fail to find effects of sex on stroke outcome and support a non-gender-biased approach to treatment [117]. Age, on the other hand, is likely to be an important factor in the diagnosis and treatment of aphasia. This points to the need for a better understanding of the relationship between language and ageing, both in the healthy and in the clinical population.

## Conclusion

Women were found to be diagnosed with aphasia following stroke more often than men. This is in direct opposition to the hypothesis that women have less lateralized language function than men. The sex difference was found to most likely be caused by age differences in the two groups at the time of stroke.

## Acknowledgements

The author has no conflict of interests to report.

## Supporting information

**S1 File. PRISMA Checklist.** Checklist for meta-analyses according to http://www.prisma-statement.org

## References

1. Benjamin EJ, Blaha MJ, Chiuve SE, Cushman M, Das SR, Deo R, et al. Heart Disease and Stroke Statistics-2017 Update: A Report From the American Heart Association. Circulation. 2017;135(10):e146–e603.

2. Ellis C, Hardy RY, Lindrooth RC, Peach RK. Rate of aphasia among stroke patients discharged from hospitals in the United States. Aphasiology. 2018;32(9):1075–86.

3. Inatomi Y, Yonehara T, Omiya S, Hashimoto Y, Hirano T, Uchino M. Aphasia during the acute phase in ischemic stroke. Cerebrovasc Dis. 2008;25(4):316–23.

4. Tsouli S, Kyritsis AP, Tsagalis G, Virvidaki E, Vemmos KN. Significance of Aphasia after First-Ever Acute Stroke: Impact on Early and Late Outcomes. Neuroepidemiology. 2009;33(2):96–102.

5. Godefroy O, Dubois C, Debachy B, Leclerc M, Kreisler A. Vascular Aphasias. Main Characteristics of Patients Hospitalized in Acute Stroke Units. Stroke. 2002;33(3):702.

6. Flowers HL, Skoretz SA, Silver FL, Rochon E, Fang J, Flamand-Roze C, et al. Poststroke Aphasia Frequency, Recovery, and Outcomes: A Systematic Review and Meta-Analysis. Arch Phys Med Rehabil. 2016;97(12):2188–201.e8.

7. Thommessen B, Thoresen GE, Bautz-Holter E, Laake K. Validity of the Aphasia Item from the Scandinavian Stroke Scale. Cerebrovasc Dis. 2002;13(3):184–6.

8. Meyer BC, Lyden PD. The modified National Institutes of Health Stroke Scale: Its Time Has Come. Int J Stroke. 2009;4(4):267–73.

9. Appelros P, Stegmayr B, Terént A. Sex differences in stroke epidemiology: a systematic review. Stroke. 2009;40(4):1082–90.

10. Berglund A, Schenck-Gustafsson K, von Euler M. Sex differences in the presentation of stroke. Maturitas. 2017;99:47–50.

11. Gall SL, Donnan G, Dewey HM, Macdonell R, Sturm J, Gilligan A, et al. Sex differences in presentation, severity, and management of stroke in a population-based study. Neurology. 2010;74(12):975.

12. Bersano A, Burgio F, Gattinoni M, Candelise L. Aphasia Burden to Hospitalised Acute Stroke Patients: Need for an Early Rehabilitation Programme. Int J Stroke. 2009;4(6):443–7.

13. Pedersen PM, Jørgensen HS, Nakayama H, Raaschou HO, Olsen TS. Aphasia in acute stroke: incidence, determinants, and recovery. Ann Neurol. 1995;38(4):659–66.

14. Miceli G, Caltagirone C, Gainotti G, Masullo C, Silveri Maria C, Villa G. Influence of age, sex, literacy and pathologic lesion on incidence, severity and type of aphasia. Acta Neurol Scand. 1981;64(5):370–82.

15. Bleses D, Vach W, Slott M, Wehberg S, Thomsen P, Madsen T, et al. The Danish Communicative Developmental Inventories: validity and main developmental trends. J Child Lang. 2008;35(03):1–19.

16. Reilly D, Neumann DL, Andrews G. Gender differences in reading and writing achievement: Evidence from the National Assessment of Educational Progress (NAEP). Am Psychol. 2018.

17. Schwartz HA, Eichstaedt JC, Kern ML, Dziurzynski L, Ramones SM, Agrawal M, et al. Personality, gender, and age in the language of social media: the open-vocabulary approach. PLoS One. 2013;8(9):e73791.

18. Wallentin M. Putative sex differences in verbal abilities and language cortex: a critical review. Brain Lang. 2009;108(3):175–83.

19. Kansaku K, Kitazawa S. Imaging studies on sex differences in the lateralization of language. Neurosci Res. 2001;41(4):333–7.

20. Shaywitz BA, Shaywitz SE, Pugh KR, Constable RT, Skudlarski P, Fulbright RK, et al. Sex differences in the functional organization of the brain for language. Nature. 1995;373(6515):607–9.

21. Baron-Cohen S, Knickmeyer RC, Belmonte MK. Sex Differences in the Brain: Implications for Explaining Autism. Science. 2005;310(5749):819–23.

22. Hausmann M. Why sex hormones matter for neuroscience: A very short review on sex, sex hormones, and functional brain asymmetries. J Neurosci Res. 2016;95(1-2):40–9.

23. Sommer IEC, Aleman A, Bouma A, Kahn RS. Do women really have more bilateral language representation than men? A meta-analysis of functional imaging studies. Brain. 2004;127(8):1845–52.

24. Hirnstein M, Hugdahl K, Hausmann M. Cognitive sex differences and hemispheric asymmetry: A critical review of 40 years of research. Laterality. 2018:1–49.

25. Moher D, Liberati A, Tetzlaff J, Altman DG, The PG. Preferred Reporting Items for Systematic Reviews and Meta-Analyses: The PRISMA Statement. PLoS Med. 2009;6(7):e1000097.

26. Rohde A, Worrall L, Godecke E, O’Halloran R, Farrell A, Massey M. Diagnosis of aphasia in stroke populations: A systematic review of language tests. PLoS One. 2018;13(3):e0194143.

27. Croquelois A, Bogousslavsky J. Stroke Aphasia: 1,500 Consecutive Cases. Cerebrovasc Dis. 2011;31(4):392–9.

28. Kyrozis A, Potagas C, Ghika A, Tsimpouris PK, Virvidaki ES, Vemmos KN. Incidence and predictors of post-stroke aphasia: The Arcadia Stroke Registry. Eur J Neurol. 2009;16(6):733–9.

29. Laska AC, Hellblom A, Murray V, Kahan T, Von Arbin M. Aphasia in acute stroke and relation to outcome. J Intern Med. 2001;249(5):413–22.

30. Roquer J, Campello AR, Gomis M. Sex Differences in First-Ever Acute Stroke. Stroke. 2003;34(7):1581.

31. Brkić E, Sinanović O, Vidović M, Smajlović D. Incidence and clinical phenomenology of aphasic disorders after stroke [Ucestalost i klinicka fenomenologija afazickih poremećaja nakon mozdanog udara.]. Med Arh. 2009;63(4):197–9.

32. Brust JC, Shafer SQ, Richter RW, Bruun B. Aphasia in acute stroke. Stroke. 1976;7(2):167.

33. Hier DB, Yoon WB, Mohr JP, Price TR, Wolf PA. Gender and Aphasia in the Stroke Data Bank. Brain Lang. 1994;47(1):155–67.

34. Engelter ST, Gostynski M, Papa S, Frei M, Born C, Ajdacic-Gross V, et al. Epidemiology of Aphasia Attributable to First Ischemic Stroke. Stroke. 2006;37(6):1379.

35. Flowers HL, Silver FL, Fang J, Rochon E, Martino R. The incidence, co-occurrence, and predictors of dysphagia, dysarthria, and aphasia after first-ever acute ischemic stroke. J Commun Disord. 2013;46(3):238–48.

36. Dickey L, Kagan A, Lindsay MP, Fang J, Rowland A, Black S. Incidence and Profile of Inpatient Stroke-Induced Aphasia in Ontario, Canada. Arch Phys Med Rehabil. 2010;91(2):196–202.

37. Kadojić D, Rostohar Bijelić B, Radanović R, Porobić M, Rimac J, Dikanović M. Aphasia in Patients with Ischemic Stroke. Acta Clin Croat. 2012;51(2):221–4.

38. Gialanella B, Bertolinelli M, Lissi M, Prometti P. Predicting outcome after stroke: the role of aphasia. Disabil Rehabil. 2011;33(2):122–9.

39. Di Carlo A, Lamassa M, Baldereschi M, Pracucci G, Basile AM, Wolfe CDA, et al. Sex Differences in the Clinical Presentation, Resource Use, and 3-Month Outcome of Acute Stroke in Europe. Stroke. 2003;34(5):1114.

40. Kelly-Hayes M, Beiser A, Kase CS, Scaramucci A, D’Agostino RB, Wolf PA. The influence of gender and age on disability following ischemic stroke: the Framingham study. J Stroke Cerebrovasc Dis. 2003;12(3):119–26.

41. Jerath NU, Reddy C, Freeman WD, Jerath AU, Brown RD. Gender Differences in Presenting Signs and Symptoms of Acute Ischemic Stroke: A Population-Based Study. Gend Med. 2011;8(5):312–9.

42. Wasserman JK, Perry JJ, Dowlatshahi D, Stotts G, Sivilotti MLA, Worster A, et al. Isolated transient aphasia at emergency presentation is associated with a high rate of cardioembolic embolism. CJEM. 2015;17(6):624–30.

43. Chang T, Gajasinghe S, Arambepola C. Prevalence of Stroke and Its Risk Factors in Urban Sri Lanka. Stroke. 2015;46(10):2965.

44. Scarpa M, Colombo A, Sorgato P, De Renzi E. The Incidence of Aphasia and Global Aphasia in Left Brain-Damaged Patients. Cortex. 1987;23(2):331–6.

45. Siirtola M, Narva EV, Siirtola T. On the occurrence and prognosis of aphasia in patients with cerebral infarction. Scand J Soc Med Suppl. 1977;14:128–33.

46. Crinion JT, Leff AP. Recovery and treatment of aphasia after stroke: functional imaging studies. Curr Opin Neurol. 2007;20(6):667–73.

47. Watila MM, Balarabe SA. Factors predicting post-stroke aphasia recovery. J Neurol Sci. 2015;352(1):12–8.

48. Plowman E, Hentz B, Ellis C. Post-stroke aphasia prognosis: a review of patient related and stroke-related factors. J Eval Clin Pract. 2011;18(3):689–94.

49. Jongbloed L. Prediction of function after stroke: a critical review. Stroke. 1986;17(4):765.

50. Ferro JM, Mariano G, Madureira S. Recovery from Aphasia and Neglect. Cerebrovasc Dis. 1999;9(suppl 5)(Suppl. 5):6–22.

51. Vuković V, Galinović I, Lovrenčić-Huzjan A, Budišić M, Demarin V. Women and Stroke: How Much do Women and Men Differ? A Review – Diagnostics, Clinical Differences, Therapy and Outcome. Coll Antropol. 2009;33(3):977–84.

52. Ellis C, Urban S. Age and aphasia: a review of presence, type, recovery and clinical outcomes. Top Stroke Rehabil. 2016;23(6):430–9.

53. Lazar RM, Boehme AK. Aphasia As a Predictor of Stroke Outcome. Curr Neurol Neurosci Rep. 2017;17(11):83.

54. Hoffmann M, Chen R. The Spectrum of Aphasia Subtypes and Etiology in Subacute Stroke. J Stroke Cerebrovasc Dis. 2013;22(8):1385–92.

55. Leśniak M, Bak T, Czepiel W, Seniów J, Członkowska A. Frequency and Prognostic Value of Cognitive Disorders in Stroke Patients. Dement Geriatr Cogn Disord. 2008;26(4):356–63.

56. Lubart E, Leibovitz A, Baumoehl Y, Klein C, Gil I, Abramovitz I, et al. Progressing stroke with neurological deterioration in a group of Israeli elderly. Arch Gerontol Geriatr. 2005;41(1):95–100.

57. Paolucci S, Antonucci G, Gialloreti LE, Traballesi M, Lubich S, Pratesi L, et al. Predicting Stroke Inpatient Rehabilitation Outcome: The Prominent Role of Neuropsychological Disorders. Eur Neurol. 1996;36(6):385–90.

58. Stegmayr B, Asplund K, Wester PO. Trends in incidence, case-fatality rate, and severity of stroke in northern Sweden, 1985-1991. Stroke. 1994;25(9):1738.

59. Stürmer T, Schlindwein G, Kleiser B, Roempp A, Brenner H. Clinical Diagnosis of Ischemic versus Hemorrhagic Stroke: Applicability of Existing Scores in the Emergency Situation and Proposal of a New Score. Neuroepidemiology. 2002;21(1):8–17.

60. Tilling K, Sterne JAC, Rudd AG, Glass TA, Wityk RJ, Wolfe CDA. A New Method for Predicting Recovery After Stroke. Stroke. 2001;32(12):2867.

61. Troisi E, Paolucci S, Silvestrini M, Matteis M, Vernieri F, Grasso MG, et al. Prognostic factors in stroke rehabilitation: the possible role of pharmacological treatment. Acta Neurol Scand. 2002;105(2):100–6.

62. Vidović M, Sinanović O, Šabaškić L, Hatičić A, Brkić E. Incidence and Types of Speech Disorders in Stroke Patients. Acta Clin Croat. 2011;50(4):491–3.

63. Lazar RM, Speizer AE, Festa JR, Krakauer JW, Marshall RS. Variability in language recovery after first-time stroke. J Neurol Neurosurg Psychiatry. 2008;79(5):530.

64. Wade DT, Hewer RL, David RM, Enderby PM. Aphasia after stroke: natural history and associated deficits. J Neurol Neurosurg Psychiatry. 1986;49(1):11.

65. Candelise L, Gattinoni M, Bersano A, Micieli G, Sterzi R, Morabito A. Stroke-unit care for acute stroke patients: an observational follow-up study. The Lancet. 2007;369(9558):299–305.

66. Hillis AE, Heidler J. Mechanisms of early aphasia recovery. Aphasiology. 2002;16(9):885–95.

67. Lloyd-Jones D, Adams RJ, Brown TM, Carnethon M, Dai S, De Simone G, et al. Heart Disease and Stroke Statistics—2010 Update. Circulation. 2010;121(7):e46.

68. Maas MB, Lev MH, Ay H, Singhal AB, Greer DM, Smith WS, et al. The Prognosis for Aphasia in Stroke. J Stroke Cerebrovasc Dis. 2012;21(5):350–7.

69. Kremer C, Perren F, Kappelin J, Selariu E, Abul-Kasim K. Prognosis of aphasia in stroke patients early after iv thrombolysis. Clin Neurol Neurosurg. 2013;115(3):289–92.

70. Pedersen PM, Vinter K, Olsen TS. Aphasia after Stroke: Type, Severity and Prognosis. Cerebrovasc Dis. 2004;17(1):35–43.

71. Hachioui HE, van de Sandt-Koenderman MWME, Dippel DWJ, Koudstaal PJ, Visch-Brink EG. The Screeling: Occurrence of Linguistic Deficits in Acute Aphasia PostStroke. J Rehabil Med. 2012;44(5):429–35.

72. Fennis TFM, Compter A, van den Broek MWC, Koudstaal PJ, Algra A, Koehler PJ. Is Isolated Aphasia a Typical Presentation of Presumed Cardioembolic Transient Ischemic Attack or Stroke? Cerebrovasc Dis. 2013;35(4):337–40.

73. Gialanella B, Santoro R, Ferlucci C. Predicting outcome after stroke: the role of basic activities of daily living predicting outcome after stroke. Eur J Phys Rehabil Med. 2013;49(3):629–37.

74. González-Fernández M, Davis C, Molitoris JJ, Newhart M, Leigh R, Hillis AE. Formal Education, Socioeconomic Status, and the Severity of Aphasia After Stroke. Arch Phys Med Rehabil. 2011;92(11):1809–13.

75. Kemper C, Koller D, Glaeske G, van den Bussche H. Mortality and Nursing Care Dependency One Year After First Ischemic Stroke: An Analysis of German Statutory Health Insurance Data. Top Stroke Rehabil. 2011;18(2):172–8.

76. Miller N, Gray WK, Howitt SC, Jusabani A, Swai M, Mugusi F, et al. Aphasia and Swallowing Problems in Subjects With Incident Stroke in Rural Northern Tanzania: A Case-Control Study. Top Stroke Rehabil. 2014;21(1):52–62.

77. Ali M, Bath PM, Lyden PD, Bernhardt J, Brady M. Representation of People with Aphasia in Randomized Controlled Trials of Acute Stroke Interventions. Int J Stroke. 2014;9(2):174–82.

78. Ojala-Oksala J, Jokinen H, Kopsi V, Lehtonen K, Luukkonen L, Paukkunen A, et al. Educational History Is an Independent Predictor of Cognitive Deficits and Long-Term Survival in Postacute Patients With Mild to Moderate Ischemic Stroke. Stroke. 2012;43(11):2931.

79. Yuan S-M, Humuruola G. Stroke of a cardiac myxoma origin. Braz J Cardiovasc Surg. 2015;30:225–34.

80. Giaquinto S, Buzzelli S, Francesco L, Lottarini A, Montenero P, In PT, et al. On the prognosis of outcome after stroke. Acta Neurol Scand. 1999;100(3):202–8.

81. Worrall LE, Hudson K, Khan A, Ryan B, Simmons-Mackie N. Determinants of Living Well With Aphasia in the First Year Poststroke: A Prospective Cohort Study. Arch Phys Med Rehabil. 2017;98(2):235–40.

82. Kauhanen ML, Korpelainen JT, Hiltunen P, Määttä R, Mononen H, Brusin E, et al. Aphasia, Depression, and Non-Verbal Cognitive Impairment in Ischaemic Stroke. Cerebrovasc Dis. 2000;10(6):455–61.

83. Bogousslavsky J, Van Melle G, Regli F. The Lausanne Stroke Registry: analysis of 1,000 consecutive patients with first stroke. Stroke. 1988;19(9):1083–92.

84. Haselbach D, Renggli A, Carda S, Croquelois A. Determinants of Neurological Functional Recovery Potential after Stroke in Young Adults. Cerebrovascular Diseases Extra. 2014;4(1):77–83.

85. Maehlum S, Roaldsen K, Kolsrud M, Dahl M. Rehabilitering etter hjerneslag [Rehabilitation after a stroke]. Tidsskrift for Den norske legeforening. 1990;110:2657–9.

86. Wade DT, Hewer RL, Wood VA. Stroke: influence of patient's sex and side of weakness on outcome. Arch Phys Med Rehabil. 1984;65(9):513–6.

87. Kelly PJ, Furie KL, Shafqat S, Rallis N, Chang Y, Stein J. Functional recovery following rehabilitation after hemorrhagic and ischemic stroke11No commercial party having a direct financial interest in the results of the research supporting this article has or will confer a benefit on the authors or on any organization with which the authors are associated. Arch Phys Med Rehabil. 2003;84(7):968–72.

88. Ween JE, Mernoff ST, Alexander MP. Recovery Rates after Stroke and Their Impact on Outcome Prediction. Neurorehabil Neural Repair. 2000;14(3):229–35.

89. Fuh JL, Wang SJ, Larson EB, Liu HC. Prevalence of Stroke in Kinmen. Stroke. 1996;27(8):1338.

90. Matsumoto N, Whisnant JP, Kurland LT, Okazaki H. Natural History of Stroke in Rochester, Minnesota, 1955 Through 1969: An Extension of a Previous Study, 1945 Through 1954. Stroke. 1973;4(1):20.

91. Safaz I, Kesikburun S, Adigüzel E, Yilmaz B. Determinants of disease-specific health related quality of life in Turkish stroke survivors. Int J Rehabil Res. 2016;39(2).

92. Kunitz SC, Gross CR, Heyman A, Kase CS, Mohr JP, Price TR, et al. The pilot Stroke Data Bank: definition, design, and data. Stroke. 1984;15(4):740–6.

93. Sarno MT, Buonaguro A, Levita E. Gender and recovery from aphasia after stroke. J Nerv Ment Dis. 1985;173(10):605–9.

94. Pickersgill MJ, Lincoln NB. Prognostic indicators and the pattern of recovery of communication in aphasic stroke patients. J Neurol Neurosurg Psychiatry. 1983;46(2):130.

95. Law J, Rush R, Pringle AM, Irving AM, Huby G, Smith M, et al. The incidence of cases of aphasia following first stroke referred to speech and language therapy services in Scotland. Aphasiology. 2009;23(10):1266–75.

96. Jodzio K, Drumm DA, Nyka WM, Lass P, Gąsecki D. The contribution of the left and right hemispheres to early recovery from aphasia: A SPECT prospective study. Neuropsychol Rehabil. 2005;15(5):588–604.

97. De Renzi E, Faglioni P, Ferrari P. The Influence of Sex and Age on the Incidence and Type of Aphasia. Cortex. 1980;16(4):627–30.

98. Basso A, Capitani E, Moraschini S. Sex Differences in Recovery from Aphasia. Cortex. 1982;18(3):469–75.

99. Oliveira FF, Damasceno BP. Short-term prognosis for speech and language in first stroke patients. Arq Neuropsiquiatr. 2009;67:849–55.

100. Sundet K. Sex differences in severity and type of aphasia1. Scand J Psychol. 1988;29(3-4):168–79.

101. Bhatnagar SC, Jain SK, Bihari M, Bansal NK, Pauranik A, Jain DC, et al. Aphasia type and aging in Hindi-speaking stroke patients. Brain Lang. 2002;83(2):353–61.

102. Eslinger PJ, Damasio AR. Age and type of aphasia in patients with stroke. J Neurol Neurosurg Psychiatry. 1981;44(5):377.

103. Kertesz A, Benke T. Sex equality in intrahemispheric language organization. Brain Lang. 1989;37(3):401–8.

104. Pizzamiglio L, Mammucari A, Razzano C. Evidence for sex differences in brain organization in recovery in aphasia. Brain Lang. 1985;25(2):213–23.

105. Kertesz A, Sheppard ANN. The Epidemiology of Aphasic and Cognitive impairment in Stroke: Age, Sex, Aphasia Type and Laterality Differences. Brain. 1981;104(1):117–28.

106. McGlone J, Kertesz A. Sex Differences in Cerebral Processing of Visuospatial Tasks. Cortex. 1973;9(3):313–20.

107. Viechtbauer W. Conducting meta-analyses in R with the metafor package. J Stat Softw. 2010;36(3):1–48.

108. Cohen J. A power primer. Psychol Bull. 1992;112(1):155–9.

109. Wallentin M, Michaelsen JLD, Rynne I, Nielsen RH. Lateralized task shift effects in Broca’s and Wernicke’s regions and in visual word form area are selective for conceptual content and reflect trial history. Neuroimage. 2014;101:276–88.

110. Wallentin M. Er der kønsforskelle i hjernens bearbejdning af sprog? Tidsskrift for sprogforskning. 2009;7:1–35.

111. Wallentin M, Skakkebæk A, Bojesen A, Fedder J, Laurberg P, Østergaard JR, et al. Klinefelter syndrome has increased brain responses to auditory stimuli and motor output, but not to visual stimuli or Stroop adaptation. Neuroimage Clin. 2016;11:239–51.

112. O'Malley KJ, Cook KF, Price MD, Wildes KR, Hurdle JF, Ashton CM. Measuring Diagnoses: ICD Code Accuracy. Health Serv Res. 2005;40(5p2):1620–39.

113. Goldstein LB. Accuracy of ICD-9-CM coding for the identification of patients with acute ischemic stroke: effect of modifier codes. Stroke. 1998;29(8):1602–4.

114. Kuznetsova A, Brockhoff PB, Christensen RHB. lmerTest Package: Tests in Linear Mixed Effects Models. J Stat Softw. 2017;82(13).

115. Lindenstrøm E, Boysen G, Waage Christiansen L, à Rogvi Hansen B, Würtzen Nielsen P. Reliability of Scandinavian Neurological Stroke Scale. Cerebrovasc Dis. 1991;1(2):103–7.

116. Hantson L, De Weerdt W, De Keyser J, Diener HC, Franke C, Palm R, et al. The European Stroke Scale. Stroke. 1994;25(11):2215–9.

117. Appelros P, Stegmayr B, Terént A. A review on sex differences in stroke treatment and outcome. Acta Neurol Scand. 2010;121(6):359–69.

